# Phosphatidylinositol (PI) Lipids Modulate the Binding of Tau Fibrils on Lipid Bilayers

**DOI:** 10.1101/2023.09.20.558589

**Authors:** Unmesh D. Chowdhury, Arnav Paul, B. L. Bhargava

**Affiliations:** School of Chemical Sciences, National Institute of Science Education & Research-Bhubaneswar, an OCC of Homi Bhabha National Institute, P.O. Jatni, Khurda, Odisha 752050,India

**Keywords:** Phosphatidylinositol lipids, clustering, tau fibrils, molecular dynamics, lipid bilayers, Alzheimer’s disease

## Abstract

Phosphatidylinositol (PI) lipids play a crucial role as a vital lipid component in cell membrane domain formation, contributing to cell signaling. In this study, we investigate the impact of PI lipids on the conformational dynamics of tubulin-associated unit (tau) fibrils through multiscale modelling. While prior experimental work by the Lecomte group has demonstrated the influence of PI lipids on the morphology and secondary structure of tau fragments, a detailed molecular understanding of the binding mechanism between tau and PI-incorporated lipids remains absent. Our molecular dynamics (MD) simulations reveal the intricate molecular mechanisms governing tau binding to PI-incorporated bilayers. Specifically, we conduct MD simulations on lipid patches containing 1-palmitoyl-2-oleoyl-sn-glycero-3-phosphocholine (PC) and 1-palmitoyl-2-oleoyl-sn-glycero-3-phosphoglycerol (PG), enabling us to explore conformational changes in the R3–R4 section of tau fibrils. Control simulations are conducted on pure lipid patches without tau fibrils, as well as on tau fibrils within bulk water. Our findings demonstrate that PI-incorporated lipids exhibit a stronger affinity for binding to tau fibrils compared to pure PC/PG lipids. All-atom simulations highlight the potential docking sites for PI headgroups at positively charged residues (Lysine, Arginine) on the tau surface. Moreover, the aggregation of PI lipids facilitates tau binding to the membrane. These results not only enhance our comprehension of the disruption of PI-incorporated bilayers, but also shed light on the stability of the tau over the PI containing bilayers.

## 1 Introduction

Neurodegenerative diseases like Alzheimer’s disease (AD), Pick’s disease, frontotemporal lobar degeneration, and progressive supranuclear palsy are all characterized by aggregation of tau protein.^1,2^ The collective term for all such diseases is tauopathies. The tau protein polymerizes tubulin into microtubules (MTs) and provides axonal support to the microtubules.^3^ Tau protein undergoes phosphorylation in post-translational modification, but sometimes hyperphosphorylation occurs when the concentration of phosphates is almost three times than that of the tau protein in a healthy brain. ^4,5^ The abnormal increase in phosphate concentration hinders the association of tau protein with microtubules and results in aggregation of tau into paired helical filaments (PHF) and neurofibrillary tangles (NFT).^6^

AD is characterized by the formation of plaques which contain aggregates of amyloid beta peptides and aggregates of tau in the form of NFTs. Research on AD is now being focussed on tau after amyloid beta targeting treatments failed in clinical trials. ^7,8^ Broadly, tau has four domains, the N-terminal domain, the proline rich domain, repeat domain region and the C-terminal domain. Tau protein in the adult human brain has four repeat domains (4R tau) and the R3 and R4 repeat domains form the core of the paired helical filament (PHF) and straight filament (SF).^9^ PHFs and SFs comprise of two protofilament cores with C-shaped subunits in different conformation, as elucidated by the cryo-EM structure by Fitzpatrick and coworkers.^10^

Studies suggest that tau interacts with the plasma membrane (PM) through its N-terminal, and the aggregation of tau is promoted by the PM in vitro. However, the mechanism is unknown.^11–14^ Membrane lipids like phosphatidylcholine (PC), cholesterol, and sphingolipid have been observed to be associated with the tau proteins. ^15,16^ Studies suggest that anionic lipid bilayers have a greater tendency to cause fibril formation in tau and A*β* proteins.^17–20^ Phosphatidylinositol 4,5-bisphosphate (PIP_2_), which is an anionic lipid, is an important lipid in the study of aggregation of tau proteins. ^20^ In an earlier work, tau and PIP_2_ binding has been reported, though structural information regarding the binding of tau over PIP_2_ lipid was absent.^21^ A more recent experimental study has shown that PIP_2_ induces aggregation of repeat domains of the tau protein and increases the random coil content in the tau protein.^22^ Phosphorylation of the phosphatidylinositol (PI) molecule at the 3, 4 and 5 position of the inositol gives rise to seven different phosphoinositide derivatives. PIP_2_, which is a major form of phosphorylated PI, is a signaling lipid, which facilitates the binding of intracellular proteins to membranes. PIP_2_ has a negatively charged inositol headgroup which can accommodate a negative charge of up to (−5), although (−3) or (−4) charge is observed in some cases. ^23^ The mammalian cell membrane has around 5 – 10% of PIs and even though it is not a major component, it serves many important biological purposes like controlling the cell death.^24^ PIP_2_ is also involved in the generation of messenger molecules, membrane trafficking, membrane protein interaction, protein – protein interactions and protein oligomerization. ^25–28^ PI lipids are found to significantly influence the dynamics and the structure of the PC membranes.^29^

Molecular dynamics simulations have paved the way to study the interaction of lipid bilayers with amyloid fibrils.^30^ MD studies also show that in zwitterionic lipids, the interaction of bilayer and protein is mostly coulombic. ^31^ However, most of the works on protein–membrane have been done on amyloid beta fibrils. MD simulation studies on tau fibril involving the lipid bilayers are still lacking. MD study of the full-length tau fibril was done by Nussinov group but without any lipid bilayer.^32^ In our earlier work, we had studied the effect of cholesterol in lipid bilayer on the interaction of tau fibril with differently charged lipid molecules.^33^ Also, we have studied the effect of tau polymorphs on the model neuronal membrane comprising of six different lipid components.^34^ But due to the lack of study on the effect of PI lipids on the interaction of tau fibril with the lipid bilayer, we carried out all-atom and coarse-grained molecular dynamics simulations taking a similar starting configuration. In the subsequent sections, PC, PG and PI are used to refer to POPC, POPG and PIP_2_ lipids, respectively. The schematics of the lipid molecules is included in the Supplementary Information (Figure S1).

## 2 Methodology and simulation details

### 2.1 Coarse-grained simulations

Martini model 2.0 and 2.2 are used to model the lipids and the proteins respectively, using CHARMM-GUI.^35^ The simulations are run using the parallelized GROMACS (version-2019). The systems were initially energy minimized and then equilibrated in six steps restraining the proteins and lipids subsequently. Electrostatics and Lennard–Jones cut-off was set to 1.1 nm with the reaction field method and potential shift Verlet method respectively. The initial velocities of the simulations were chosen from a Maxwell distribution at 310K. The temperature and the pressure for the production run was kept constant using v-rescale thermostat and Parrinello–Rahman barostat respectively. The coupling constants of v-rescale thermostat and Parrinello–Rahman barostat were 1 ps and 12 ps respectively. The production runs were performed for 10 *µ*s with a time step of 20 fs. Electrostatic interaction was modelled using the reaction field method using the dielectric constant of 15.

### 2.2 All-atom simulations

The initial structures of the protein were taken from the pdb ids 5O3T and 5O3L. The proteins have been modelled using the CHARMM-36m force field. CHARMM-36m is shown to perform best for the PHF6 motif of the tau peptides among a series of force fields.^36,37^ The simulations were performed using the GROMACS MD code (version-5.1.4).^38^ The fibril was initially placed above the lipid bilayers containing PIP_2_ in the ratio of 9:1 lipid:PIP_2_. The water model TIP3P is used along with an additional 0.15(M) KCl to maintain the physiological conditions. The lipid molecules and PIP_2_ were modelled using the CHARMM-36 parameters developed by Klauda et al.^39,40^ Lipids were subjected to six steps of equilibration (by applying restraints on proteins and lipids subsequently) after the initial energy minimization following the CHARMM-GUI protocol.^35,41^ The final production runs were performed at the temperature of 310K using the Nosé–Hoover thermostat with a coupling constant of 1.0 ps. Production runs were performed in the NPT ensemble for 200 ns using a time step of 2 fs. Long ranged electrostatics were modelled using the Particle Mesh Ewald (PME) with a cut-off of 1.2 nm.^42^ Force switch function was used to model the long range interactions. The bonds with the hydrogen atoms were restrained using the LINCS algorithm.^43^The tau fibril/membrane simulations have been replicated twice to increase the conformational sampling.

For the water box simulations, the polymorphs (SF and PHF) of the tau fibrils were packed in a cubic box with edge length of 1.4 – 1.5 nm. Pure lipid systems without the protein have been modelled with 200 lipid molecules in the ratio of 9:1 of the PC/PG and PI lipids. The lipids were packed symmetrically with 100 lipids in both the upper and the lower leaflet. The bilayer models were generated using the CHARMM-GUI Membrane Builder. 200 ns of trajectory was generated from the production runs in the NPT ensemble at 310 K and 1 bar. All the analyses were done using *gmx* tools and using in-house scripts invoking MDAnalysis python module using the entire length of the trajectory.^44^ The PI clusters are calculated using DBSCAN (Density-based spatial clustering of applications with noise) algorithm from *sklearn.cluster* library. The distance cutoff of 1.5 nm is chosen to characterize the clusters.

### 2.3 System Description

We have simulated symmetrical lipid bilayers consisting of PC/PG + PI lipids. The tau fibrils are placed above the lipid bilayers consisting of PI lipids in accordance with the earlier studies.^33^ To increase the configurational sampling we have replicated the simulations twice. The results are calculated as the average of these two simulations. Further details of the simulated systems are given in Table 1. The lipids are having a composition of PC/PG and PI lipids in the ratio of 9:1. The structures of straight filament and paired helical filament are shown in Figure 1(a)-(b) respectively. The fibril–lipid starting configuration is given in Figure 1(c).

**Figure 1:**
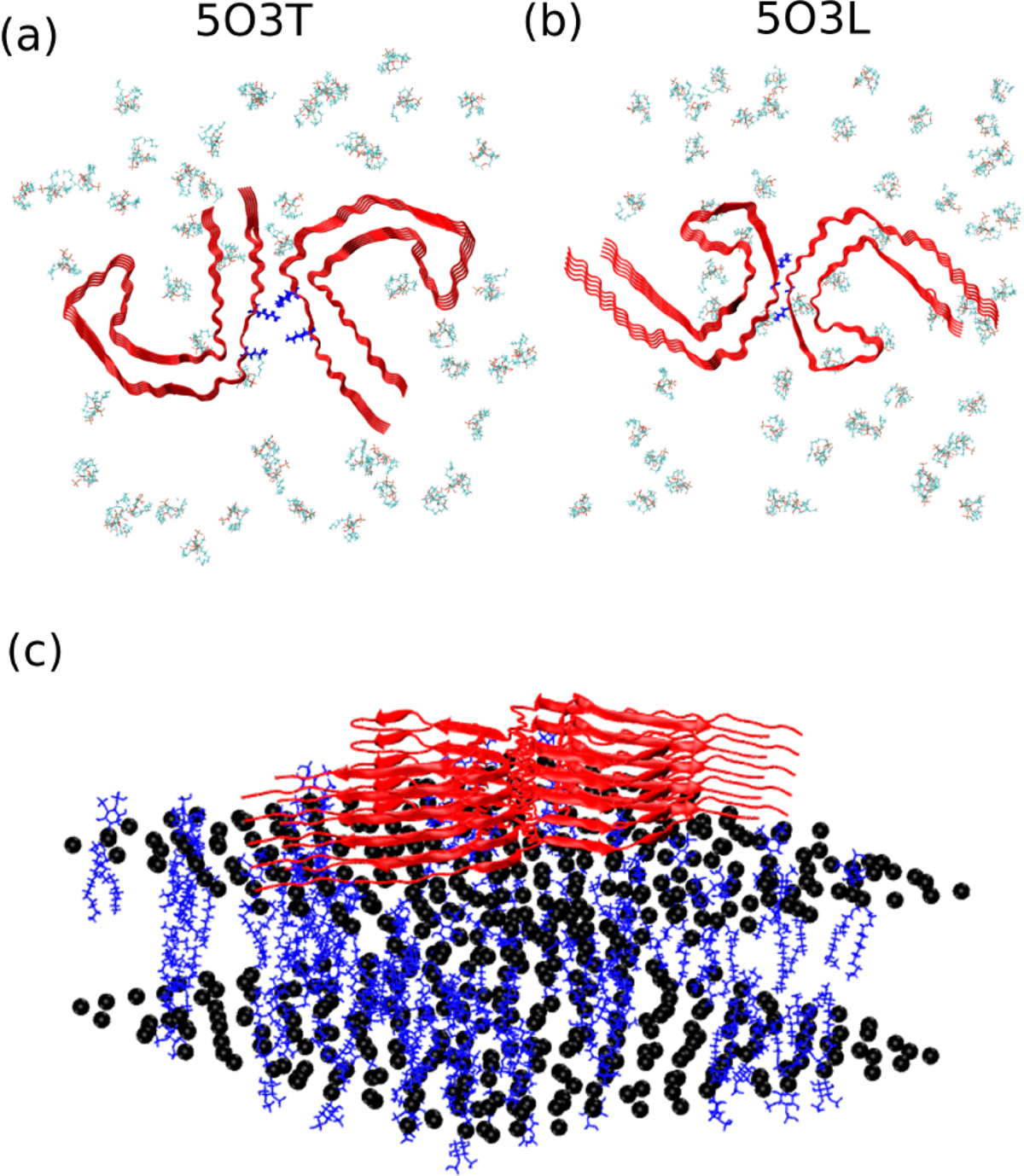
(a) and (b) are the straight filament (SF) and paired helical filament (PHF) corresponding to the PDB ids 5O3T and 5O3L respectively, along with the PI lipids as viewed over the top. The PI lipids are shown in lines representations in VMD. Residues K317, K321 of SF and the residues G334, Q336 of PHF are shown in blue. (c) is the initial configuration of the systems with the fibril over the bilayer. The PI lipids are shown as blue sticks and the phosphorus headgroups of the PC lipids are shown in black.

**Table 1:** Summary of the simulations reported in this paper.

| System description | Number of lipids | Production run | Total number of atoms/beads (for coarse grained simulations) |
| --- | --- | --- | --- |
| All-atom (protein and lipids) |  |  |  |
| PC+PI (SF) | 510 | 200 ns × 2 | 225094 |
| PC+PI (PHF) | 600 | 200 ns × 2 | 374443 |
| PG+PI (SF) | 510 | 200 ns × 2 | 275756 |
| PG+PI (PHF) | 600 | 200 ns × 2 | 178524 |
| Pure Lipids |  |  |  |
| PC+PI | 510 | 400 ns | 63086 |
| PG+PI | 510 | 400 ns | 58754 |
| Tau-protein (water box) |  |  |  |
| SF | — | 500 ns | 332578 |
| PHF | — | 500 ns | 393490 |
| Coarse-grained (protein and lipids) |  |  |  |
| PC+PI (SF) | 510 | 10 μs | 54972 |
| PC+PI (PHF) | 510 | 10 μs | 56060 |
| PG+PI (SF) | 510 | 10 μs | 51463 |
| PG+PI (PHF) | 510 | 10 μs | 51539 |
| Pure Lipids |  |  |  |
| PC+PI | 510 | 9 μs | 38813 |
| PG+PI | 510 | 9 μs | 44190 |

## 3 Results and Discussions

### 3.1 Tau Fibril Stability

The root mean square fluctuations (RMSFs) of the C-*α* atoms of the tau fibrils are given in Figure 2(a). The root mean square deviations (RMSDs) of the fibrils for all the systems are included in the Supplementary Information (Figure S2–S5). All simulations show convergence in RMSD values. The density distribution of the PI lipids projected over the bilayer plane is included in Supplementary Information (Figure S6). RMSF provides information about the mobility and flexibility of atoms or residues within a protein. High RMSF values indicate more flexibility and greater deviation from the average position, suggesting dynamic regions or regions undergoing conformational changes. Conversely, low RMSF values suggest more rigid or less mobile regions. If the distribution of the C shape angle of coverage is narrow and centered around a specific value, it suggests that the tau fibril adopts a stable and well-defined C-shaped conformation. This indicates that the fibril structure is relatively rigid and maintains its characteristic shape throughout the simulation. If the distribution of the C shape angle is broad, it suggests that the tau fibril exhibits conformational flexibility. The PHF residues exhibit greater stability compared to the SF structures. The difference between the RMSF values of tau in the PC+PI and PG+PI is greater for the SF structure compared to the PHF structure. Also, the residues are more flexible in the presence of the PC+PI lipids in the SF structure. This is due to the lower affinity of the tau fibril to bind with the bilayer in PC+PI systems, which is discussed in a later section.

**Figure 2:**
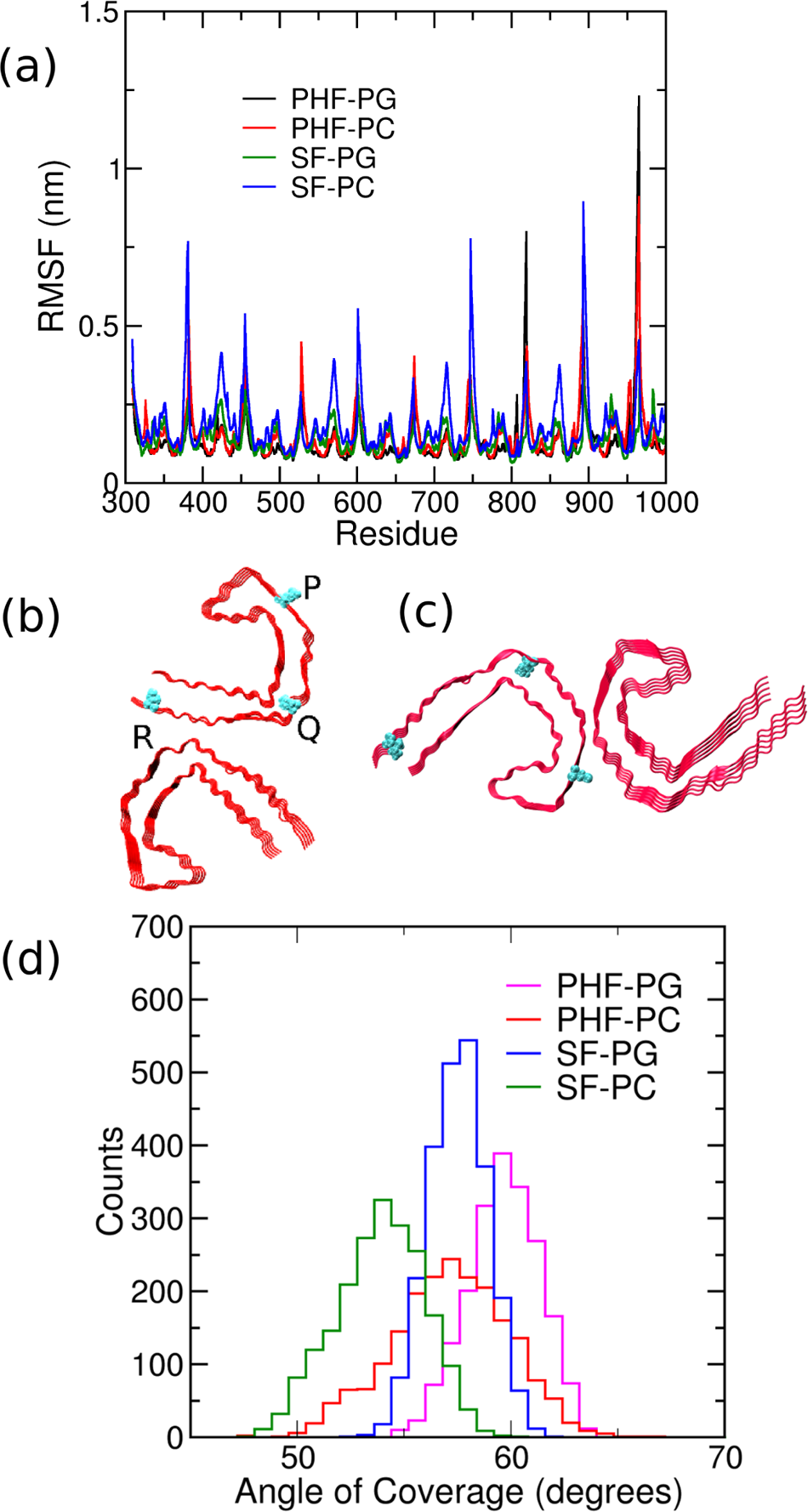
(a) RMSF of the C-*α* residues of the tau fibrils. (b)-(c) are the angle of coverage PQR used in the discussion. (d) The distribution of the angle of coverage in different systems. PHF/SF–PC/PG stands for the PHF or SF structures in POPC/POPG + PIP_2_ bilayers.

The angle of coverage is defined by the residues as shown in Figure 2(b)-(c). The distribution of the angle of coverage is shown in Figure 2(d). The increase in the angle of coverage is correlated with the fluctuations of the tau residues in case of the PHF fibril in PG+PI systems. The most probable angle of coverage is minimum (53°) in case of SF structure in PC+PI system, which increases to a maximum value of 59° in the PHF structure in PG+PI system. The corresponding ‘angle of coverage’ for the PHF and the SF structures in the cryo-EM structures are 58° and 57° respectively. We infer that the increase in the angle of coverage is due to the higher affinity of the fibril to bind with PI incorporated lipid bilayer.

### 3.2 Bilayer Properties

#### 3.2.1 Area per lipids and Bilayer thickness

The bilayer thickness refers to the distance between the surface of the two leaflets of the lipid bilayer. It is measured as the distance between the phosphate groups of the two lipid layers. Bilayer thickness is affected by several factors, including the length and saturation of the lipid tails, the size and shape of the lipid headgroup, and the presence of cholesterol or other membrane proteins.^45,46^ The thickness of the bilayer can influence the diffusion of small molecules and the conformational dynamics of membrane proteins.^47^

The area per lipid refers to the amount of area that each lipid occupies in the bilayer. It is measured as the total area of the lipid bilayer divided by the number of lipids in the bilayer. Area per lipid is influenced by the size and shape of the lipid headgroup and the length and saturation of the lipid tails. The area per lipid can affect the packing and fluidity of the membrane, which can impact the diffusion of small molecules and the activity of membrane proteins.^46,48,49^ In general, changes in bilayer thickness and area per lipid can alter the physical properties of the lipid bilayer, which can have important biological consequences.

To elucidate the fibril lipid interactions, we have looked at the changes in the bilayer properties. The area per lipids (APL) and the bilayer thickness in the systems shed light on the effect of the fibrils on the bilayers. The bilayer thickness is calculated by taking the average distance between the phosphate groups on either layer. The APL for the systems along with the bilayer thickness are shown in box plots in Supplementary Information (Figure S7). The mean values of the APL and the bilayer thickness are given in Table 2. The bilayer thickness and APL describe the membrane deformations induced by the tau binding. The tau binding leads to membrane thinning and also leads to the decrease of APL compared to the control systems comprising of pure lipid bilayers. This is consistent with the earlier reports concerning the amyloid *β* and also the tau fibrils.^33,50,51^

**Table 2:** Mean values and standard deviations of the area per lipid (nm^2^) and bilayer thickness (nm) of tau incorporated systems and control systems. SF/PHF–PC/PG refers to the SF or PHF structure with PC+PI and PG+PI lipids respectively.

|  | Area Per Lipid (nm<sup>2</sup>) | Bilayer Thickness (nm) |
| --- | --- | --- |
| Pure PI-PC | 0.64 ± 0.01 | 3.89 ± 0.06 |
| Pure PI-PG | 0.69 ± 0.01 | 3.70 ± 0.05 |
| PHF-PC | 0.55 ± 0.01 | 3.83 ± 0.05 |
| PHF-PG | 0.60 ± 0.01 | 3.67 ± 0.03 |
| SF-PC | 0.59 ± 0.01 | 3.86 ± 0.04 |
| SF-PG | 0.65 ± 0.01 | 3.53 ± 0.02 |

The two-dimensional bilayer thickness in various bilayers and the control system without the tau fibril are shown in Figure 3. The thickness is projected over the bilayer plane and is shown as a colorbar. We find that the inclusion of tau fibrils induces local deformations in the membrane surface. The local thinning of the bilayer is most important to decipher the binding of tau fibrils. In case of the PHF structure in PC+PI lipid, the bilayer thickness is large almost throughout the bilayer. In comparison, the SF structure in PC+PI lipid shows two distinct islands of lower thickness separated by larger thickness. The PHF structure in the PG+PI lipids show a comparatively thinner bilayer surface than the PC+PI system, while in the case of SF structure in the PG+PI lipids, there are certain small regions of greater thickness in comparison to the SF structure in the PC+PI lipids. Overall, we find that the PG+PI bilayer in presence of PHF structure show the largest decrease in the bilayer thickness in the tau-incorporated membrane. The PC+PI bilayer in the presence of PHF structure exhibits the largest membrane thickness signifying reduced binding over the membrane.

**Figure 3:**
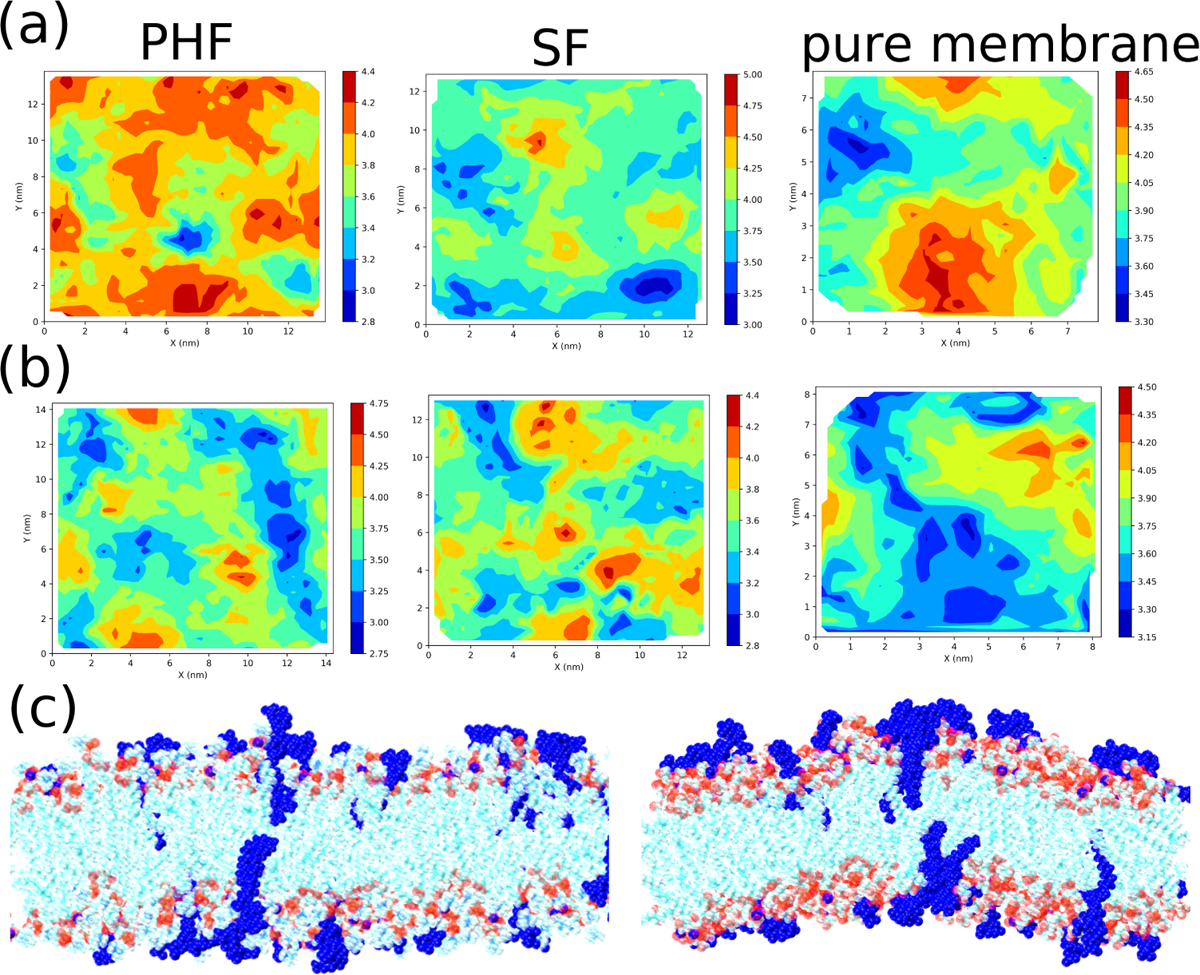
(a)-(b) are the bilayer thickness for the systems projected over the bilayer plane for the PC+PI and PG+PI systems respectively. The bilayer thickness (nm) is shown in a colorbar. (c) shows the snapshots of the tau incorporated PC+PI (left) and PG+PI (right) systems.

#### 3.2.2 Order Parameter

Order parameter (S*_CH_*) of the lipid tails signify the time-averaged C–H bond angle, with respect to the bilayer normal. The Order Parameter of the acyl chains is defined as follows

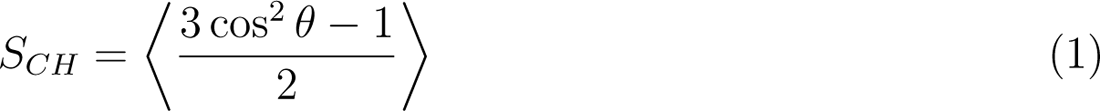

The S*_CH_* value of 1 corresponds to completely ordered acyl chains, and S*_CH_* value *→* 0 signifies randomly oriented acyl chains. The S*_CH_* values of the sn-1 and the sn-2 chains in all the systems are shown in Figure 4 for PC and PG lipids. For both chains, the first seven carbon atoms show larger S*_CH_* values than atoms in the membrane core region, suggesting higher membrane fluidities near the lipid core. The highest values for the acyl chains are found in the pure PC–PI systems followed by SF–PC system. The least ordering of the acyl chains are found in the SF fibril with PG+PI lipids. Incorporation of PHF and SF tau fibrils decreases the lipid ordering as found from the earlier studies.^19,33^ Yet, there is only negligible change (approximately 2%) in the PC+PI systems in going from the pure membrane to the tau incorporated membrane. On the other hand, there is a significant decrease in the order parameter in going from the pure PG+PI lipids to the fibril incorporated lipid membrane with the highest decrease in the PHF structure (nearly 16%). Hence, we conclude that the PHF fibril perturbs the PG+PI more effectively, than the SF fibril. Also, the negatively charged PG lipids are more affected by the tau fibrils compared to the PC lipids.

**Figure 4:**
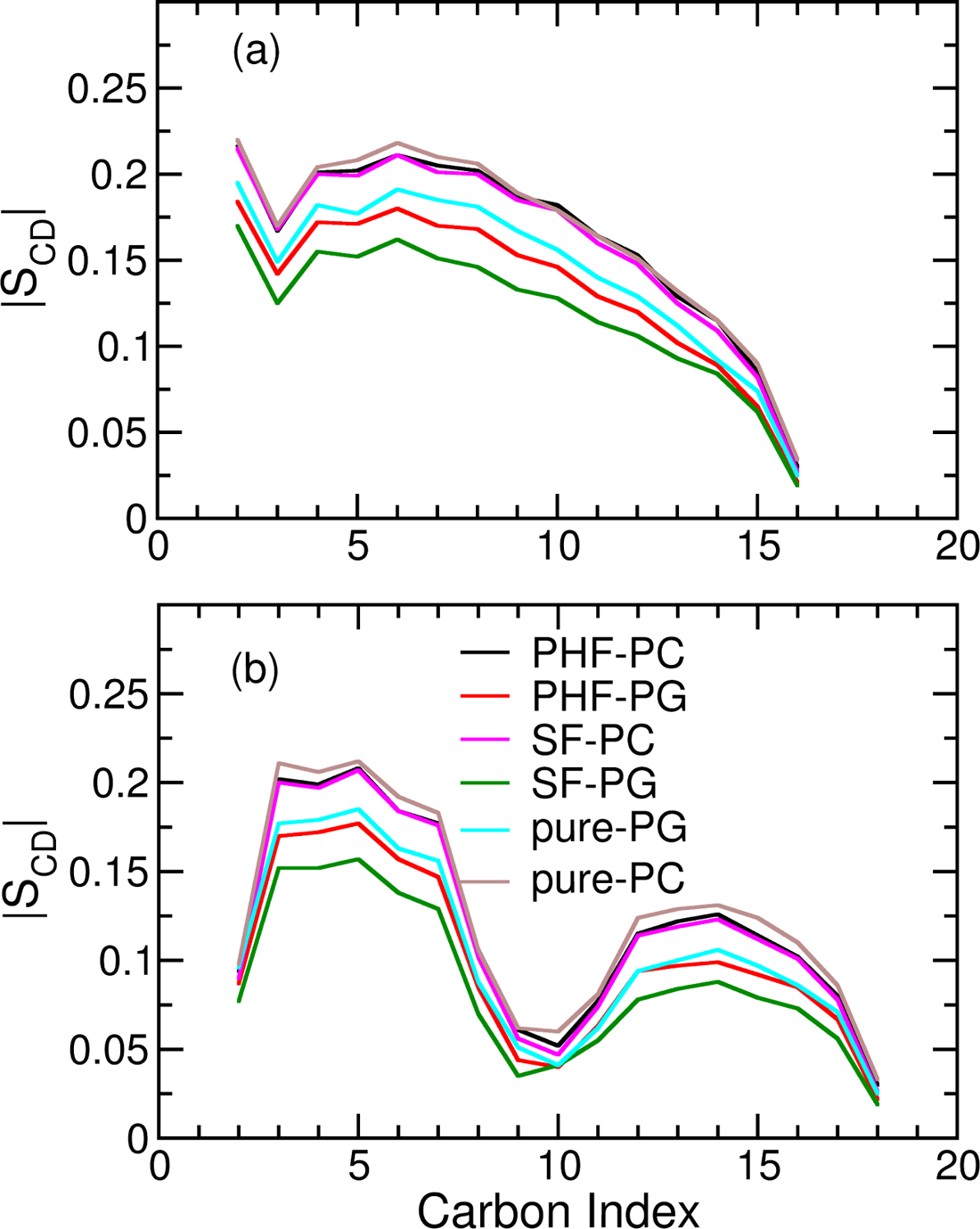
(a)-(b) are the order parameters of the sn-1 and sn-2 lipid tails of PC/PG, respectively.

#### 3.2.3 Membrane Roughness

Membrane roughness is defined by the following equation^52^

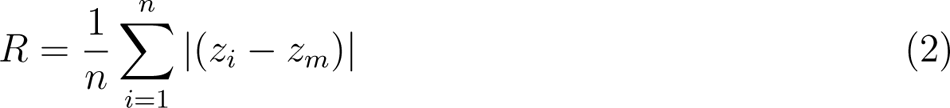

where n is the total number of reference atoms, z*_i_* is the z-coordinate of reference atom i, and z*_m_* is the mean z-position of reference atoms. The values of the membrane roughness are given in Table 3. The membrane roughness is calculated from the average z-position of the lipid which describes the vertical fluctuations of the lipid molecules in the bilayers. We have taken the phosphorus head group of PG or PC lipid as the reference atom in calculating the roughness.

**Table 3:**
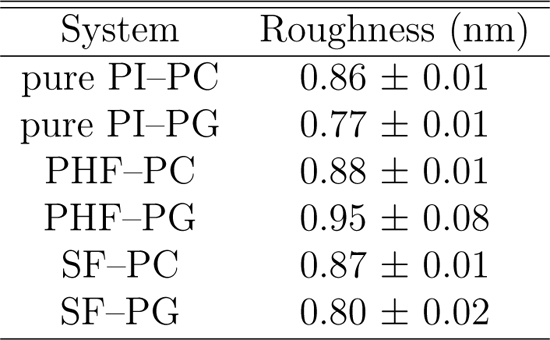
Membrane Roughness (nm) of tau incorporated systems and control systems.

| System | Roughness (nm) |
| --- | --- |
| pure PI-PC | $0.86 \pm 0.01$ |
| pure PI-PG | $0.77 \pm 0.01$ |
| PHF-PC | $0.88 \pm 0.01$ |
| PHF-PG | $0.95 \pm 0.08$ |
| SF-PC | $0.87 \pm 0.01$ |
| SF-PG | $0.80 \pm 0.02$ |

Membrane roughness can modulate the availability, accessibility, and organization of lipid molecules, thereby impacting the binding affinity and localization of proteins that interact with PI lipids. A snapshot of the PI incorporated bilayer without the tau, illustrating the membrane roughness is shown in Supplementary Information (Figure S8). Leonenko’s group showed experimentally that the membrane roughness of the diseased membrane model in the presence of the A*β* changes over time.^53^ The higher membrane roughness correlated with increasing accumulation of A*β* aggregates over the membrane. In our study, we found that the presence of tau fibrils in the system increase the membrane roughness compared to the pure PC+PI and PG+PI systems. Interaction with the PHF structure leads to the increase in the membrane roughness much more than the SF structure.

#### 3.2.4 Lipid Interdigitation

The interdigitation of the membrane lipids is calculated through the degree of interdigitation between the acyl tails.^54^ Lipid interdigitation is calculated from the width of the overlapping regions (w*_ρ_*). Earlier studies have found that membrane binding with the proteins leads to the thinning of the membrane which in turn leads to the mass overlap of the acyl chains in the opposing leaflets.^55,56^ The values of the interdigitation metrics for all the systems are given in Table 4.

**Table 4:** The interdigitation metrics for all the systems.

| System | $w_\rho$ (nm) |
| --- | --- |
| pure PI-PC | $0.51 \pm 0.03$ |
| pure PI-PG | $0.58 \pm 0.02$ |
| PHF-PC | $0.59 \pm 0.03$ |
| PHF-PG | $0.78 \pm 0.08$ |
| SF-PC | $0.60 \pm 0.05$ |
| SF-PG | $0.88 \pm 0.05$ |

We find that the decrease in the bilayer thickness correlates with the increasing interdigitation with the interaction of the tau fibrils. The interdigitation is more prominent in PG+PI in presence of SF fibrils. The interdigitation of the pure PG+PI lipids increases by 0.20 nm with the inclusion of the PHF, whereas it increases by 0.30 nm in presence of SF fibril. The change in interdigitation values of the acyl chains is minimal in the PC+PI lipids upon inclusion of tau polymorphs. Thus, we conclude that lipid interdigitation and bilayer thinning occur simultaneously with the inclusion of tau polymorphs, and the change in interdigitation is most prominent in the SF system comprising PG+PI lipids.

### 3.3 Tau bilayer interaction

#### 3.3.1 Number of Contacts

To model longer timescales of the tau-membrane interaction, we calculated the total number of contacts between the membrane and the tau structures from the coarse grained simulations. The cutoff for the contacts is chosen to be 0.7 nm. The total number of contacts between the membrane and the tau structure is shown in Figure 5(a). The distances between the center of mass of the tau fibril and the lipid bilayer is included in the Supplementary Information (Figure S9). The distances are nearly the same for all the systems. The change in the number of contacts is marginal in both the PC+PI and PG+PI systems. PHF in PC+PI bilayer shows a lower number of contacts compared to the PG+PI. On the other hand, the changes in the number of contacts in SF with PC+PI and PG+PI are minimal (nearly 1%). Hence, it is observed that the PHF structure shows the most stable binding with PG+PI bilayer. Overall, we have found a large increase (nearly 84%) in the number of contacts between the tau and the PG+PI lipids compared to the pure PG lipids. We have used the coarse grained simulation trajectory of tau with pure PG from our earlier work. ^33^ The clustering of the PI lipids is a necessary phenomenon responsible for cell signaling that is critical for cell health. ^57–60^ The clustering is mediated by the influence of cations like Ca^2+^, which is also observed in physiological conditions. ^61^ Based on the coarse grained simulations studies of our earlier report,^33^ we have compared the number of tau – lipid contacts in the SF fibril in the pure PG bilayer with that in the PG–PI bilayer as shown in 5(b). The pure membranes without the tau fibrils are also simulated for 9 *µ*s as the control for the coarse grained simulations. The snapshots of the pure membrane simulations without the tau fibrils are given in the Supplementary Information (Figure S10–S11). The clustering of the PI lipids in the control simulations are calculated using the DBSCAN algorithm which is also included in the Supplementary Information (Figure S12). The number of contacts in the systems comprising PC and PC–PI lipids are shown in Supplementary Information (Figure S13). We can infer from the coarse grained simulations that PI lipids facilitate the tau fibril binding with the bilayer. Even without the tau fibrils, the PI lipids show clustering as evident from our control simulations. Experimental and computational studies showed the tight binding between PI(4,5)P2 and K-Ras4b protein which is a vital protein for the cellular signaling through receptor tyrosine kinases.^62^ KRAS4b is more tightly bound overall with PIP2+DMPS compared to the pure DMPS.^62,63^ We also observe clustering of the PI lipids in the bilayer during the course of the simulations, which is shown in Figure 5(c)-(d). The number of PI lipids found in the clustering is shown in Figure 5(e). We predict a possible correlation between the clustering of the PI lipids and the increasing number of contacts between the tau fibril and the PI containing bilayers. Earlier studies from the coarse grained modeling has shown that the electrostatic interaction between the PI head group and the positively charged residues of Bin-Amphiphysin-Rvs (BAR) protein induce membrane binding.^64^ The clustering of the PI lipids is also found to increase the ability of the BAR protein to bind to the membrane changing the membrane curvature. In our coarse grained simulation of the PI incorporated lipids we also found the PI induced membrane curvature, which is shown in the Supplementary Information (Figure S14). Hence, we envisage that the PI lipids induce membrane curvature through the binding with the tau fibrils which in-turn is initiated through the PI clustering. Our results also agrees with the earlier reports of the mechanism of PI binding with several other proteins.^63,64^

**Figure 5:**
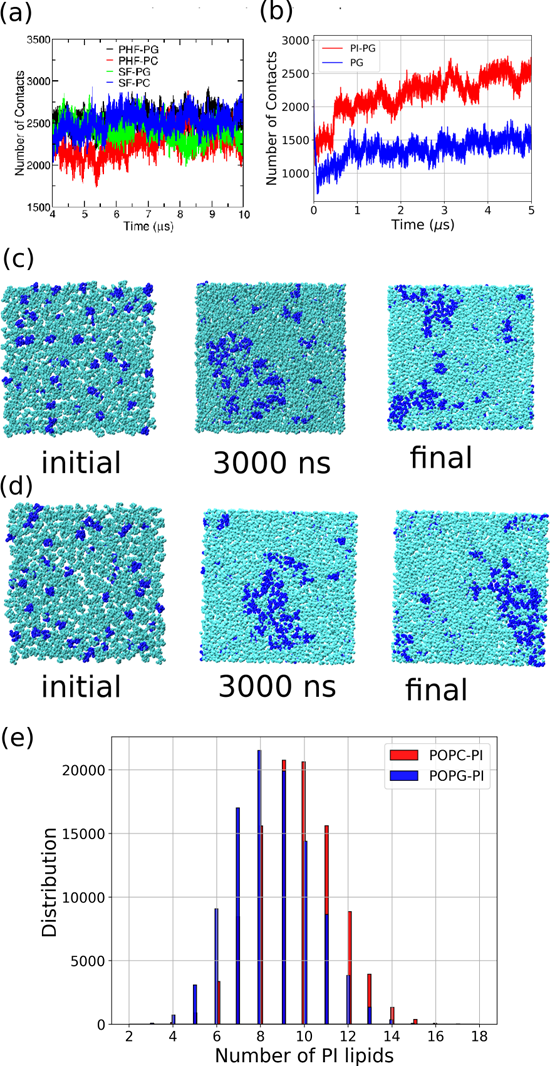
(a) The number of contacts between the tau fibrils and the bilayer from the coarsegrained simulations shown from 4–10 *µ*s. (b) Number of contacts between the tau fibrils and the bilayers for PG–PI and pure PG systems. The number of contacts for SF tau with PG systems are included from the coarse grained simulations from our earlier work. ^33^ (c)-(d) are the snaphots of the bilayer at various timesteps from the coarse grained simulations in PC and PG systems, respectively. The PI headgroups are shown as blue VDW spheres, the PC/PG beads are shown in cyan. (e) The histograms of the number of PI lipids forming clusters which is obtained from the coarse grained simulations in PHF system.

#### 3.3.2 Per Residue Contacts

To analyse the fibril–lipid contacts, we have calculated the per residue contacts of the tau polymorphs. The per residue contacts for the SF and PHF structures are shown in Figure 6. The contacts are calculated using a cutoff value of 1.0 nm from tau residue to the lipid, using the all atom trajectories. This will help us in knowing which specific amino acid residues of the tau fibrils come close to the lipid molecules, providing valuable insights into the molecular interactions that may be relevant for understanding the tau mediated pathogenesis.

**Figure 6:**
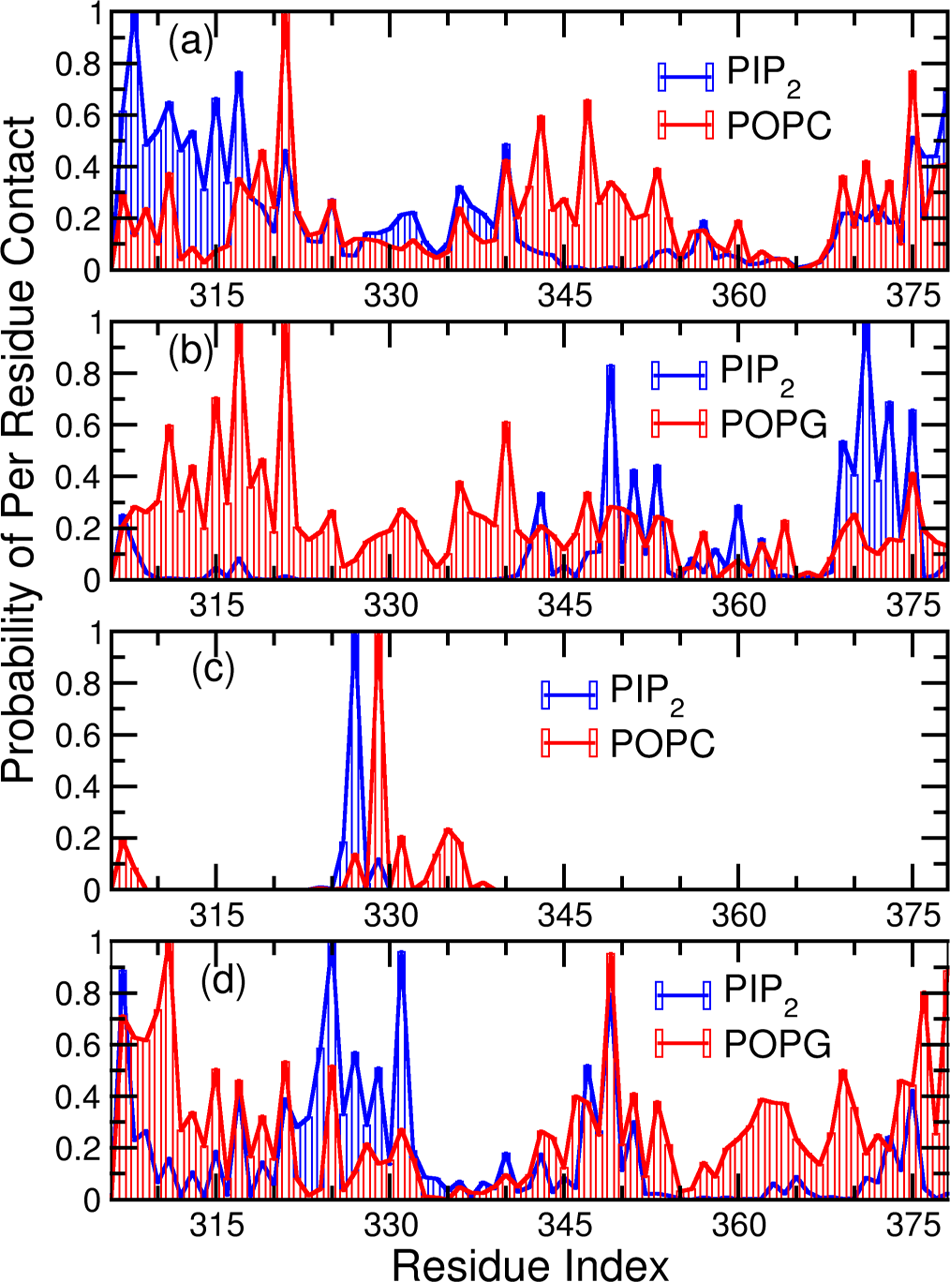
(a)-(b) are the per residue contacts for the PHF systems and (c)-(d) are the per residue contacts for the SF systems.

The PHF structures show more number of contacts with both the PC and PI lipids in comparison to the SF structures. In the case of the PHF structures, N-terminal region shows a higher number of contacts with the PI lipids, especially between the I308 and K317 residues. The higher number of PI lipid contacts leads to the concomitant decrease of the PC contacts. The PC contacts are highest for the K321 residue. Similarly, for the PHF with PG+PI lipids, the peaks for POPG lipids are seen at K317, K321 residues and I371 for the PI lipids. The least number of contacts are seen for the SF structure with PC+PI lipids with the peaks at 327–328 residues for the PI and PC lipids respectively.

#### 3.3.3 Tau dimer interface

The tau R3–R4 structure is stabilized by certain key residues in the interface. The polymorphism arises from the difference in the dimer interface though the core residues of the R3–R4 fragment are the same. The important amino acid interactions stabilizing the dimer interface are shown in Figure 7. The distance profiles of the interface residues are shown in the Supplementary Information (Figure S15). In the PHF interface, the important interactions are salt-bridge contacts between K331–E338 and the interaction between G334 residues in the adjacent chains. In the SF structure, the dimer interaction is between L315–K321 and K317–K321 residues.

**Figure 7:**
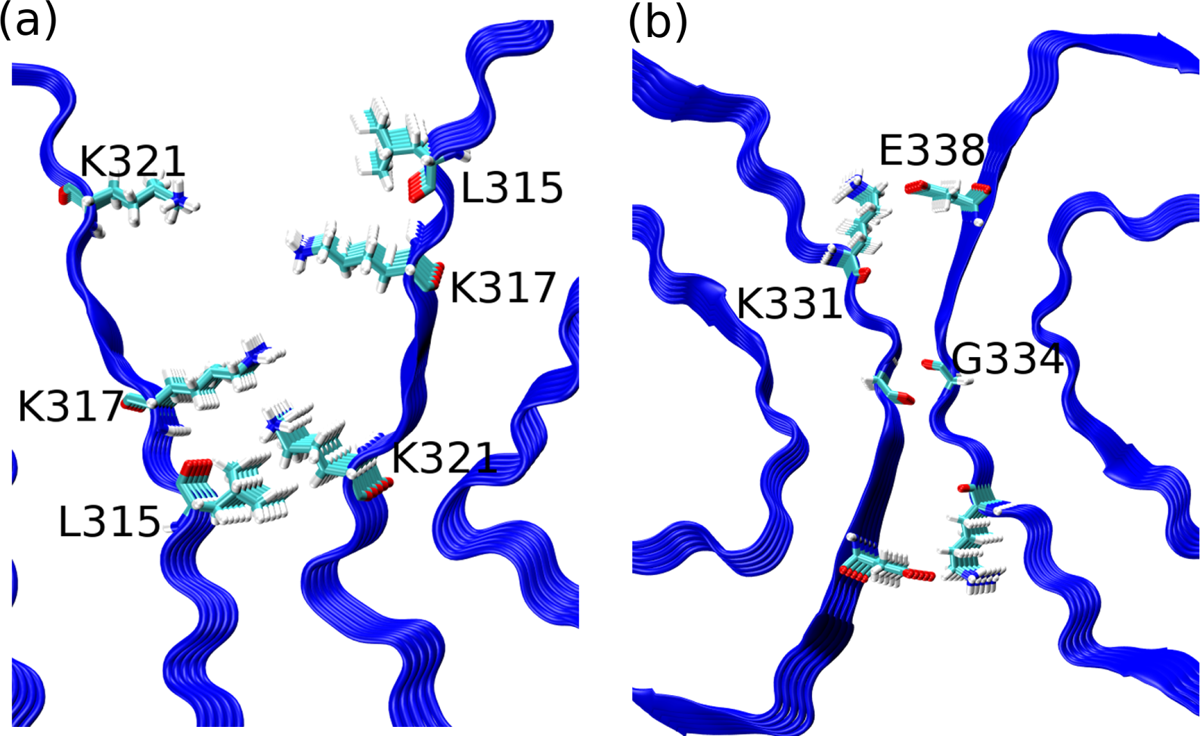
(a)-(b) are the snapshots of the tau fibril interface showing the key residues of the SF and PHF structure respectively.

In the PHF structure, the G334–G334 distance is 0.599 nm in the cryo-EM structure which decreases in all the systems. The average distance between the G334–G334 residues in PG+PI and PC+PI is nearly 0.440 nm. The salt bridge distance between the NH^+^ and COO*^−^* groups in the cryo-EM structure is 0.693 nm and in the water medium it is 0.371 nm. Hence the G334–G334 and salt bridge distance decreases in both the water medium and the PI incorporated lipids.

In the SF structure, L315–K321 residues have a distance of 0.935 nm in the cryo-EM structure and 0.783 nm in the water medium. This distance changes to 0.839 nm and 0.729 nm in the PC+PI and PG+PI systems. The distance between the K317–K321 residues changes to 1.316 nm and 1.119 nm in the PC+PI and PG+PI lipids respectively. The K317– K321 distance in the water medium is 1.260 nm and the corresponding distance in the cryo EM structure is 1.213 nm. Thus, the cryo EM distances for the interface residues changes in both the water medium and in the PI incorporated membranes. We see the tighter packing of the tau dimers in the PI incorporated bilayers in the timescales of our simulations.

#### 3.3.4 Secondary structure content

The secondary structure content of the tau polymorphs shed light on the changes in the secondary structure upon binding with the PI incorporated lipids. The earlier studies have shown that the composition of the lipid bilayers have an effect on the secondary structure content of the tau fibrils and the interaction of the tau fibrils with the lipid bilayers lead to the concomitant loss of the *β*-sheet structures.^33,65^ The cryo-EM structure of the tau R3–R4 fragments span the residues V306–K311 in *β*1, V313–C322 in *β*2, N327–K331 in *β*3, Q336–S341 in *β*4, K343–K347 in *β*5, R349–I357 in *β*6, S356–V363 in *β*7 and N368–F378 in *β*8. The per residue propensity of the tau residues is shown in Figure 8(a)-(d). The total number of the *β*-sheet residues is shown in a violin plot in Figure 8(e).

**Figure 8:**
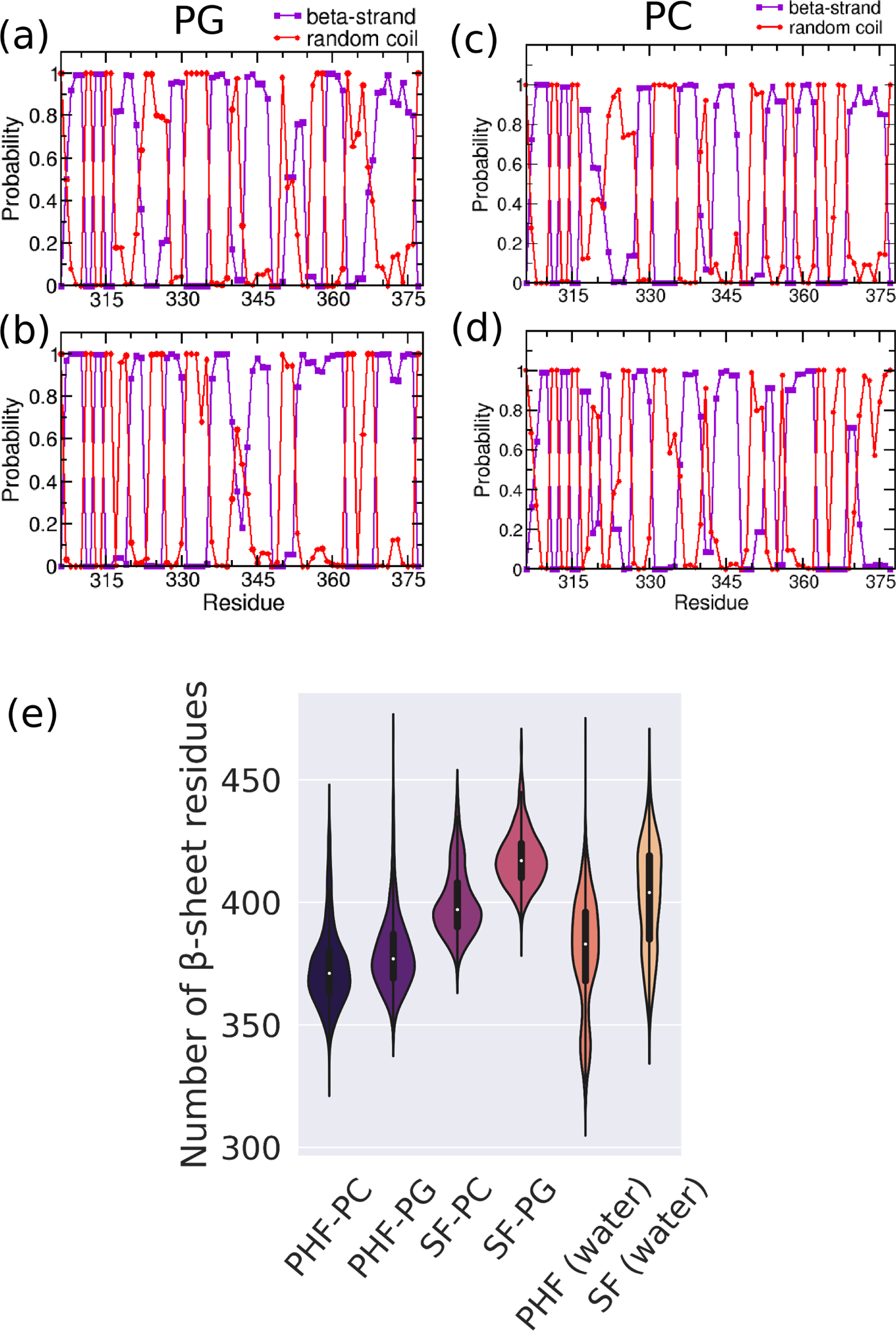
(a)-(b) are the per residue propensities in tau polymorphs PHF and SF structures respectively in the PI incorporated PG lipids, and (c)-(d) are the per residue propensities in the PI incorporated PC lipids (e) is the violin plot of the total number of *β*-sheet residues in the four systems. PHF/SF – PC/PG stands for the PHF or SF structures in PC/PG+PI bilayers.

The secondary structure content of the tau fibrils varies considerably in the presence of the PI incorporated bilayers. The PHF structures show only a marginal change in the per-residue propensity in going from PC+PI to PG+PI. The C-terminals show considerable change in the proportion of the *β*-strand and the coil conformation in the PI incorporated PC and PG lipids. Also, the *β*6 region between R349–I357 and *β*7 region between S356–V363 show the most notable changes. In the PC+PI lipids the *β*3 region and the C-terminal region show most difference in the *β*-sheet content. In terms of the total number of *β*-sheet residues, we find that the negatively charged PG+PI lipids preserve most of the *β*-sheet region in the SF structure. The least number of *β*-sheet residues is found in the PHF structure with the PC+PI lipids. In comparison with the tau-PI systems, the total number of beta-sheet residues in the water medium is 379*±*22 for the PHF fibrils and 401*±*21 in the case of SF structures. Hence, we conclude that the tau fibril interacting with the negatively charged PG with PI lipids has the most number of *β*-sheet residues. The interaction of the tau fibrils with the lipid bilayers lead to the change in number of *β*-sheet residues when compared to the tau fibrils in the bulk water.

#### 3.3.5 Bilayer Fibril Distance

To characterize the binding of the tau fibrils over the membrane bilayers, we have computed the *z*-coordinate center of the mass distance between the tau fibrils and the bilayer. The *z*-coordinate represents the vertical axis perpendicular to the surface of the bilayer, and it is used to describe the position of molecules along this axis. This describes how far the tau fibrils are from the lipid bilayer, providing valuable information about the interaction between them. This bilayer fibril approach distance is shown in Figure 9(a). The final snapshots of all the systems are shown in Figure 9(b)-(e). The tau fibril in the PG+PI lipids show a closer approach to the bilayer compared to the PC+PI lipids. The distance is less for the SF–PG system than that for the PHF–PG system. On the contrary, the SF–PC systems move farthest from the bilayer among all the systems. Thus, tau fibrils in the presence of the PG–PI lipids show the most effective binding over the membrane. To further corroborate this fact, the final snapshots are also included in Figure 9. As found in the case of the SF–PC system, the final snapshot shows the fibril moving away from the bilayer.

**Figure 9:**
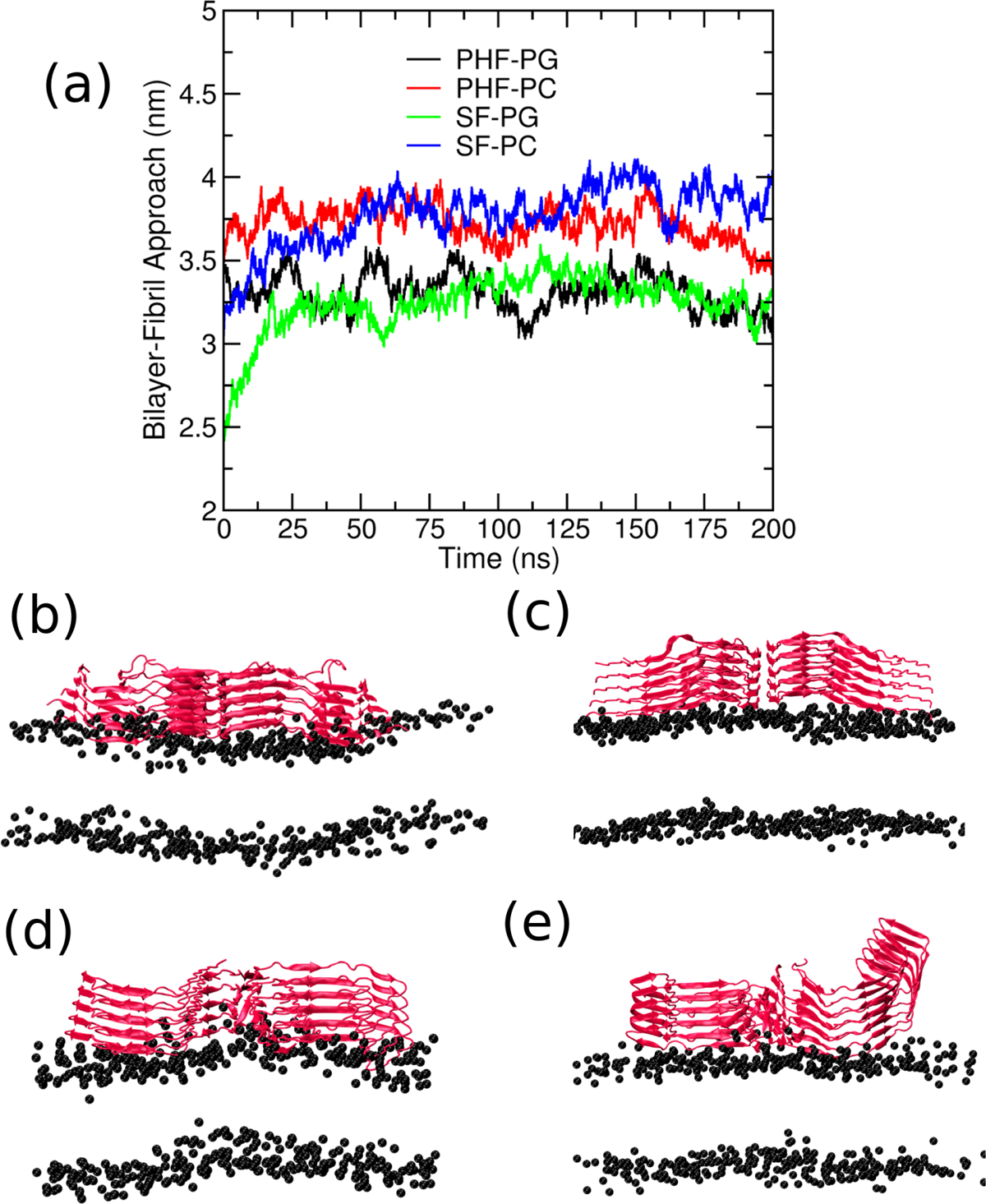
(a) The bilayer approach distance of the fibril and the bilayer along the z-direction. (b)-(c) are the final snapshots for the PHF–PG and PHF–PC systems and (d)-(e) are the final snapshots of the SF–PG and SF–PC systems.

#### 3.3.6 Binding specificity of PI lipids with the tau fibrils

The PI lipids cluster spontaneously in the lipid membrane which is responsible for cell signalling and calcium influx in the cell.^60^ The calcium ions bridge between the negatively charged phosphate groups in the PI lipids and facilitate the aggregation.^58^ It is known that the tau binding on the lipid membranes are mediated by the positively charged residues of the tau surface and the negatively charged lipid headgroups. Hence, we looked into the possible docking mechanism of the tau fibril over the PI domains on the bilayer.^34,66^ Figure 10 shows the snapshots of the interaction of tau filament (PHF) with PI lipids at different timeframes. R349 and K353 residues are shown along with a neighbouring PI lipid. The PI lipid is bound to R349 residue initially up to 55 ns. Once the PI lipid is unbound, it binds to the neighbouring K353. This binding of K353 with the PI lipid has longer residence time and it lasts nearly 160 ns. The distance between the -NH_3_^+^ of R/K residues and the -PO_4_^3^*^−^* of PI lipid is also included in Figure 10(e).

**Figure 10:**
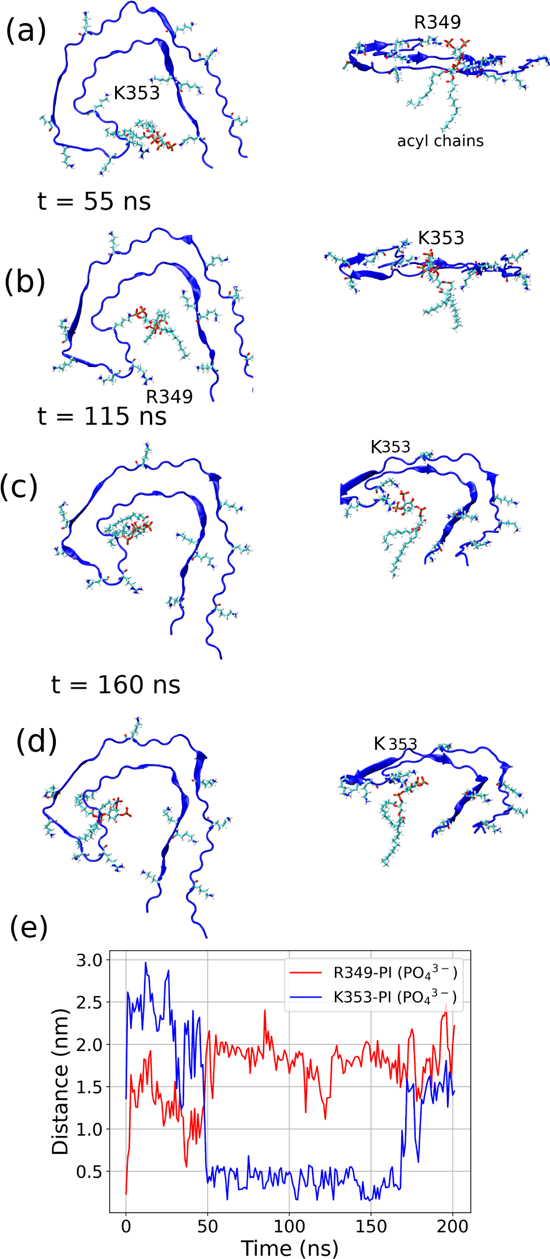
The snapshots of tau filament (PHF) interacting with the PI lipids at four different timeframes. Only the first chain of the tau fibril interacting with the lipid is shown for clarity. The positively charged residues (R, K, H) and the interacting PI lipid are shown in licorice representation. The second column is the snapshot viewed from the xz plane.

The docking mechanism of the tau residues in the PC–PI bilayer is shown in the Supplementary Information (Figure S16,S17). It is found that there is no strong binding between PI and the positively charged residues. This is also evident from the distance v/s time plot shown in the Supplementary Information (Figure S18). The phosphate groups in the inositol ring of PI lipid interacts with both the residues K343 and K347. Eventually the PI lipid moves far away from the positively charged residues on the tau surface. Thus, in the presence of PC, the residence time of the negatively charged phosphate groups and the positively charged K343, K347 is lower when compared to the PG–PI system. We postulate that the PI binding is mediated by the negatively charged PG lipids, the presence of which makes the positively charged tau residues move closer to the PI headgroups. On the contrary, the initial interaction of the tau residues with the PC lipids are less as the PC is zwitterionic. The positively charged residues of the tau are hence far away from the negatively charged phosphate groups of PI lipids which causes reduced affinity of tau binding in PC–PI lipids. Corbin et al. proposed a two-step search process based on their investigation of the GRP1 PH domain and PI lipid.^29,67^ Along with the PI lipids the bilayer comprised of anionic phosphatidylserine (PS) lipids. Initially, the PH domain engages in transient and weak electrostatic interactions with the anionic lipids (PS/PI) in the background. This interaction enhances the protein’s residency time on the membrane surface and enables a two-dimensional search for PI. Once PI is located, the PH domain binds to it specifically and with a high affinity. The first step involves proteins being attracted to the membrane surface by anionic lipids (PS and PI) through nonspecific electrostatic interactions. This interaction reduces the search dimension from three to two, facilitating efficient protein diffusion without tight binding. In the second step, proteins may encounter their target PIs in a distinct region during two dimensional diffusion, where the PI are bound to adjacent anionic lipids (PS and PI) through hydrogen bonding. Alternatively, proteins can locate target PI through orientational changes when the PI are bonded to the PS/PI in the same region. Finally, proteins bind to PIPs through specific interactions with the phosphate groups of the inositol ring or through stronger nonspecific electrostatic interactions with basic residues in the protein.

## 4 Conclusion

The interaction of tubulin-associated unit (tau) with the cellular membrane appears to be an important factor in the development of tau-related pathologies, and further research is needed to fully understand the mechanisms underlying this interaction and its contribution to disease. Utilizing tip-enhanced Raman spectroscopy, the presence of phosphatidylinositol (PI) phospholipid within tau filaments is studied earlier, offering a perspective into protein secondary structure at a nanoscale (less than 10 nm) level of detail. Yet, the atomic resolution of the tau association with the PI lipids are not yet studied. MD simulation studies of the tau polymorphs (PHF and SF) in presence of the model membrane composed of PC/PG and PI lipids in the ratio of 9:1 have been carried out to probe the interaction of phosphatidylinositol (PI) with tau fibrils. Integrating findings from both the coarse grained and atomistic simulation approaches allowed us to develop a comprehensive model of the tau-pathology interaction, which is crucial for understanding the mechanisms underlying various tauopathies and designing potential therapeutic interventions. PHF structures are found to be more stable than the SF in the PI incorporated PC and PG lipids. The tau binding over the PI containing bilayer causes membrane deformation which is evident from the order parameter, membrane interdigitation and membrane roughness. Overall, the tau fibrils are more tightly bound in the PI incorporated lipids than the pure PG/PC lipids as shown from the coarse grained simulations. We also speculate the specific binding modes with the positively charged residues, Arg, Lys, His of the tau interface and the negatively charged PI lipid headgroups. The aggregation of the PI headgroup leads to the concomitant increase of the number of contacts between the tau fibrils and the PI incorporated lipids. Though simulations done in the timescales described cannot capture all modes of interaction and adsorption of tau over the membrane, our studies provide insights into the interaction of PI with tau fibrils in bilayers comprising of PI incorporated PC and PG lipids. Multiple starting configurations with different orientations of the tau are required to capture other modes of interaction. Also, metadynamics and umbrella sampling can be done taking suitable collective variables. Overall, our results shed light on the tau binding to the phosphatidylinositol incorporated bilayers, which are known to play an important role in the cell signalling.

## Acknowledgement

The authors gratefully acknowledge NISER Bhubaneswar for the computational resources.

## Supporting Information Available

The supplementary information comprises of the schematics of the lipid molecules comprising the bilayers used in our study (Figure S1), root mean square deviations (RMSDs) for all the systems (Figure S2–S5), two dimensional density plots for all the systems (Figure S6), box plots for the area per lipid and the bilayer thickness (Figure S7), snapshot of the bilayer surface illustrating the roughness (Figure S8), distances between the center of mass of the tau fibril and the lipid bilayer from CG simulations (Figure S9), the snapshots of the initial and the final configurations of the pure lipids (Figure S10, S11), the histograms of the number of PI lipids forming clusters (Figure S12), number of contacts between the tau fibrils and the bilayers (Figure S13), the snapshot of the final configuration and the mean curvature (Figure S14), distance profiles (Figure S15), snapshots illustrating the interaction of Lys 353 and phosphate group of the inositol ring (Figure S16, S17), distance between phosphate group of inositol and Lys 343 and Lys 347 residues (Figure S18).

