## Supplementary information for "Phosphatidylinositol (PI) Lipids Modulate the Binding of Tau Fibrils on Lipid Bilayers"

S14 (a) Snapshot of the final configuration of the coarse grained simulations of the tau and the PG–PI lipids. PI lipids are shown in blue using VDW representation. Note the anchoring of the tau fibril by PI lipids. (b)-(c) are the membrane curvature for the PC+PI and PG+PI system respectively. The mean curvature is shown in a colorbar. Negative value of the membrane curvature represents concave surface and positive value of membrane curvature represents convex surface. The mean membrane curvature is calculated using the MDAnalysis python library. . . . . . . . . 13
S15 (a)-(b) are the L315–K321 distance and K317–K321 distance in the SF structure. (c)-(d) are the K331–E338 distance and G334–G334 distance in the PHF structure. The residues are shown in the main manuscript. . . . . . . . . . . . . . . . . . . . . . . . . . 14
S16 The snapshot of the part of the fibril showing the interaction between the positively charged Lys 353 and the inositol group of the PI lipid. The water molecules, lipid molecules and the other chains of the tau fibrils are omitted for clarity. . . . . . . . . . . 15
S17 The snapshots of the part of the fibril showing the positive charged residues of tau interacting with the PI inositol groups in the PC–PI systems. The water molecules, the other chains of the tau fibril and the PC lipids are omitted for clarity. . . . . . . . . . . 16
S18 The distance between the phosphate group of the inositol ring of the PI lipid and the residues Lys 343 and Lys 347. . . . . . . . . 17

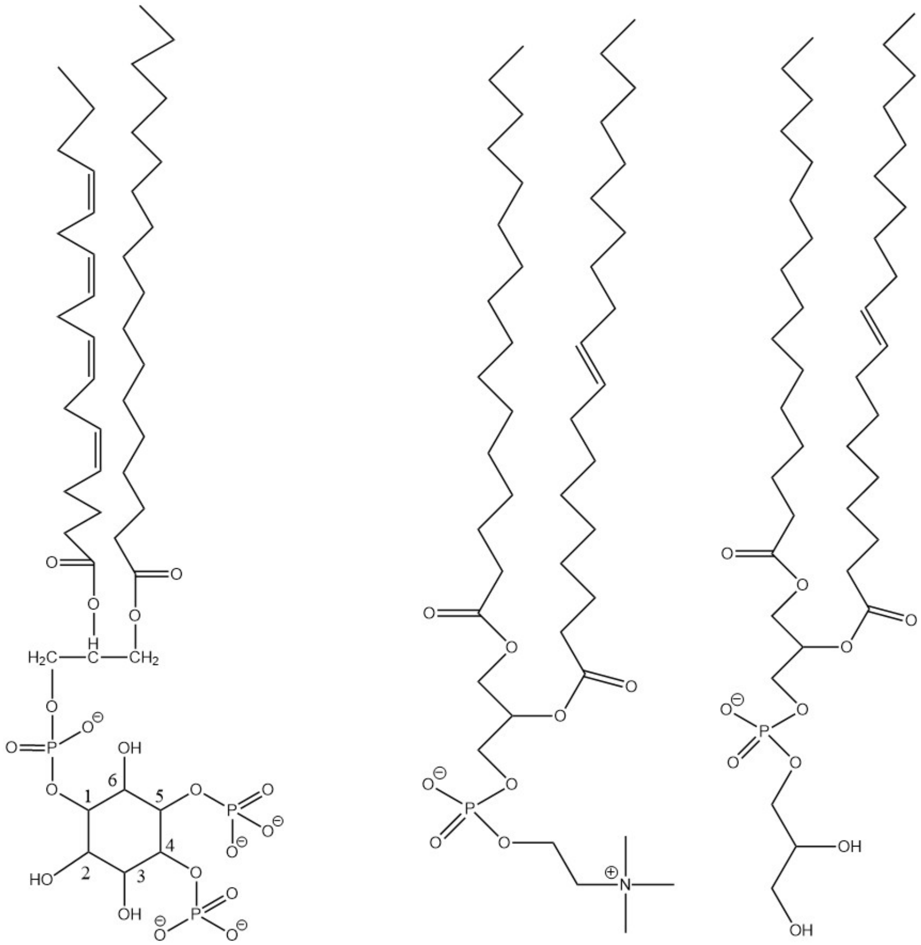

PIP2 POPC POPG

Figure S1: The schematics of lipid molecules PIP2 (PI), POPC (PC) and POPG (PG) used in our model bilayers.

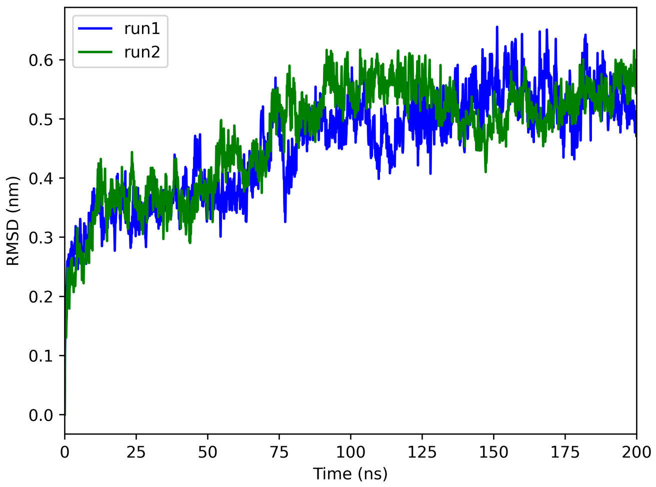

Figure S2: RMSD of the fibril in two independent simulations of SF – PC+PI system.

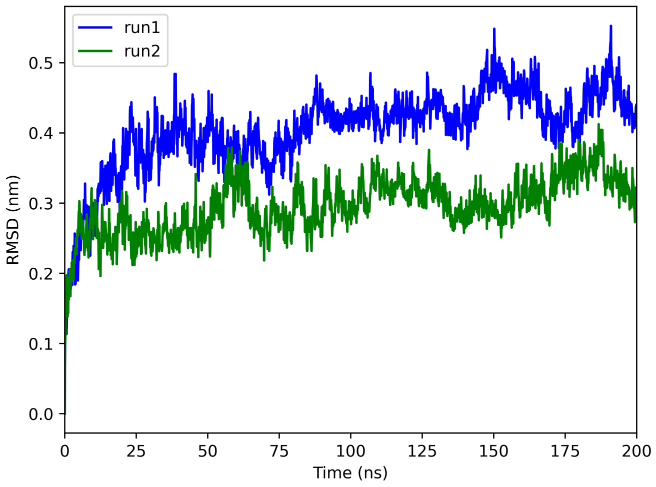

Figure S3: RMSD of the fibril in two independent simulations of PHF – PC+PI system.

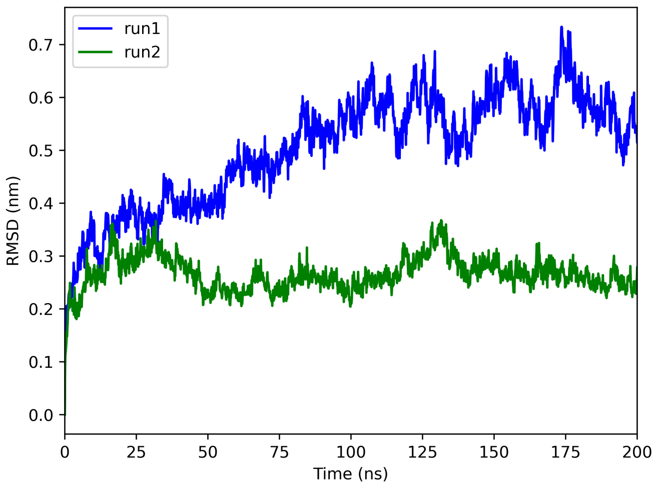

Figure S4: RMSD of the fibril in two independent simulations of SF – PG+PI system.

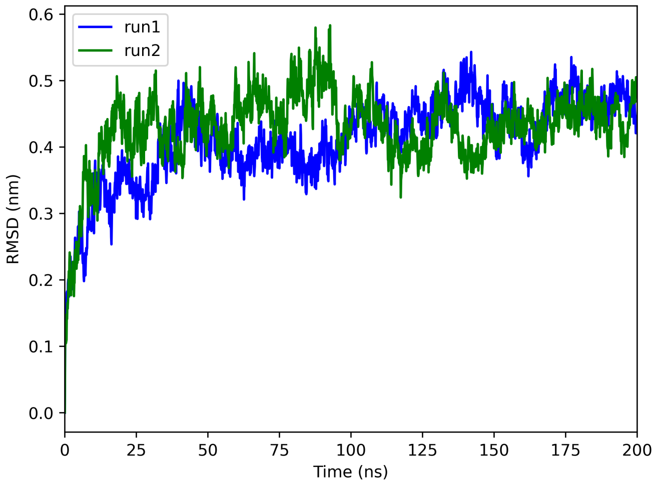

Figure S5: RMSD of the fibril in two independent simulations of PHF – PG+PI system.

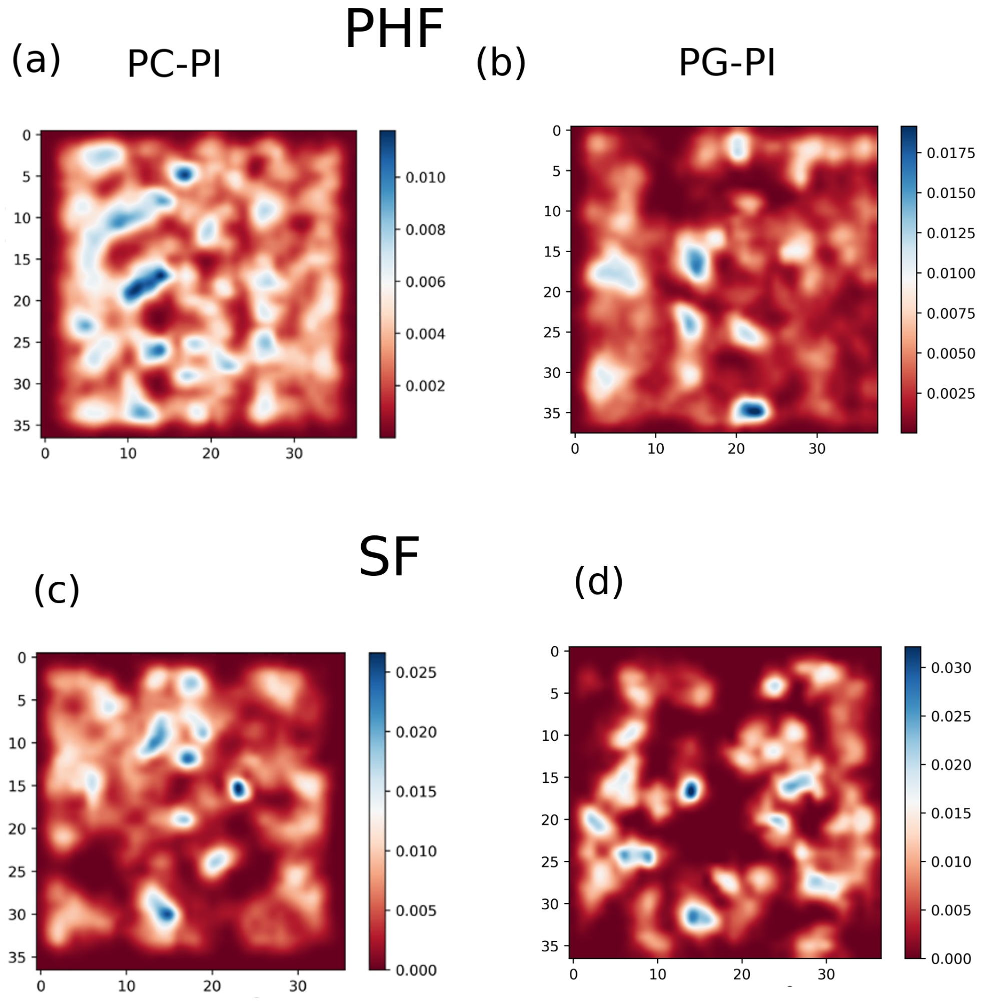

Figure S6: Density distribution of the PI lipids projected over the bilayer plane. The relative density is shown in a colorbar.

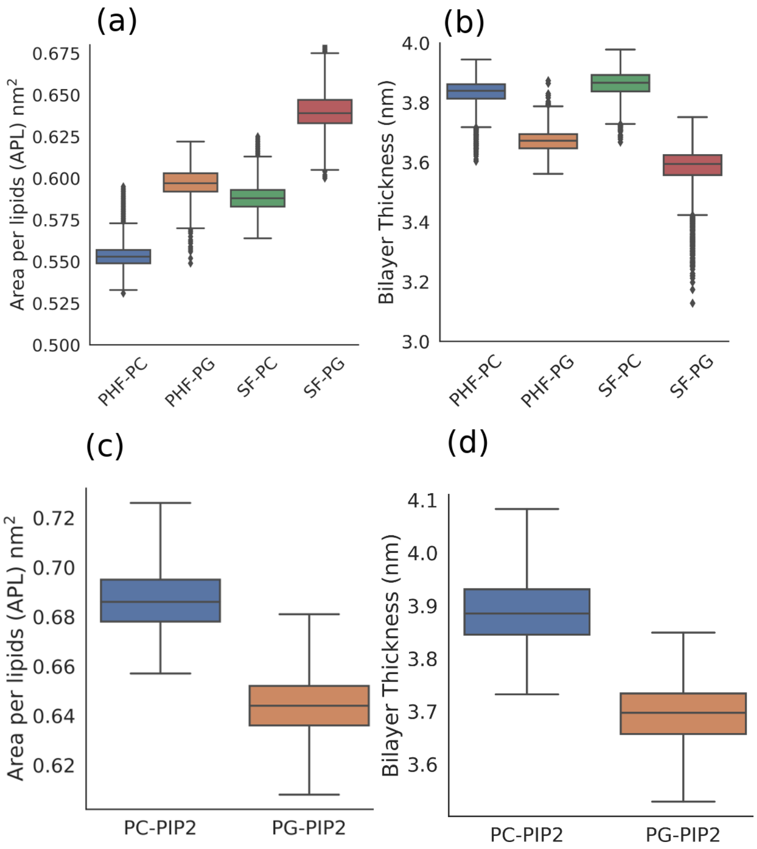

Figure S7: (a)-(b) are the box plots for the area per lipid and bilayer thickness in the four systems averaged over the entire trajectory. The APL and bilayer thickness for the control systems without the tau fibrils are shown as box plots in (c) and (d) respectively.

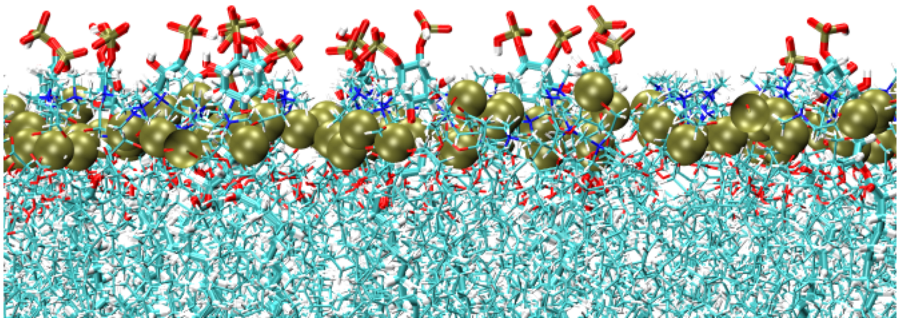

Figure S8: Snapshot of the surface of the membrane with PC and PI lipids showing the membrane roughness, from all atom modeling. The tail lipids are shown in cyan, phosphorus head groups are shown in tan as VDW spheres, inositol molecules in the PI lipids are shown in licorice representation.

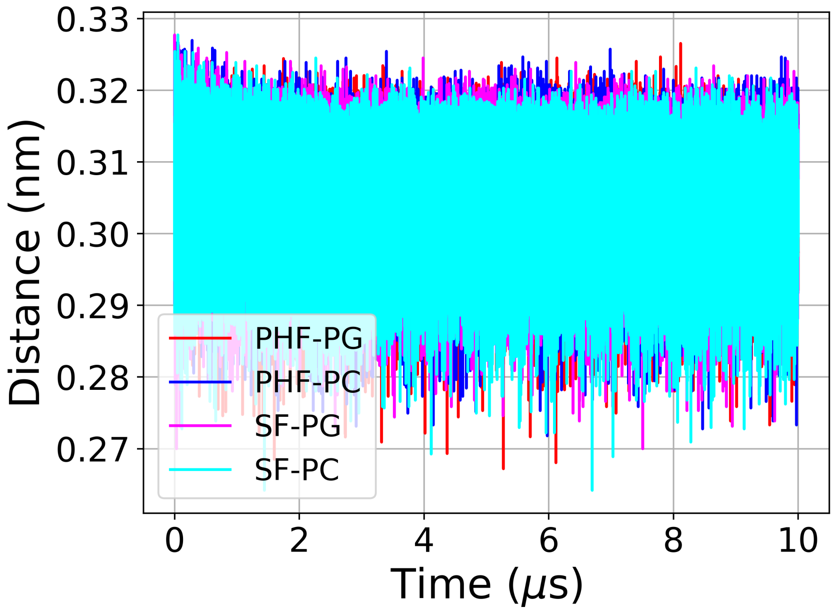

Figure S9: Distance between the center of mass of the tau fibril and lipid bilayer derived from the coarse grained simulations.

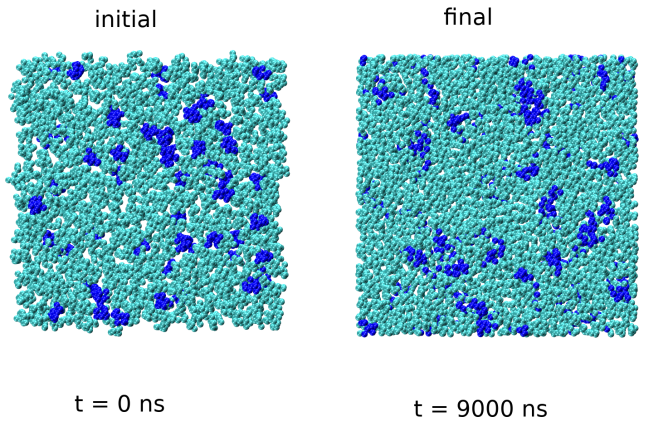

Figure S10: Snapshots of the initial and the final configuration of the pure PG-PI lipids from the CG simulations. The PG and PI lipids are shown in cyan and blue respectively.

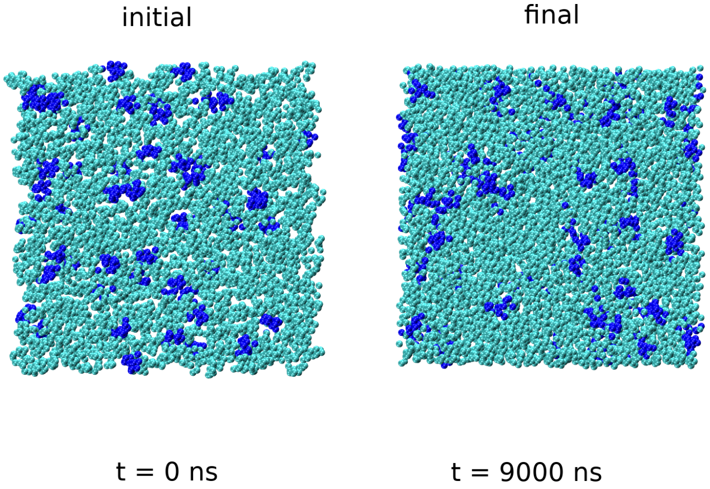

Figure S11: Snapshots of the initial and the final configuration of the pure PC-PI lipids from the CG simulations. The PC and PI lipids are shown in cyan and blue respectively.

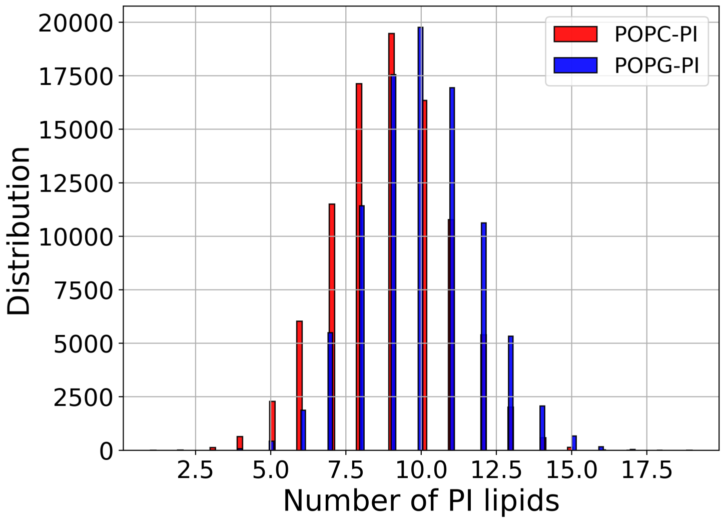

Figure S12: The histograms of the number of PI lipids forming clusters, obtained from the coarse grained simulations in the pure PC/PG and PI lipids. The clusters are obtained through the DBSCAN algorithm.

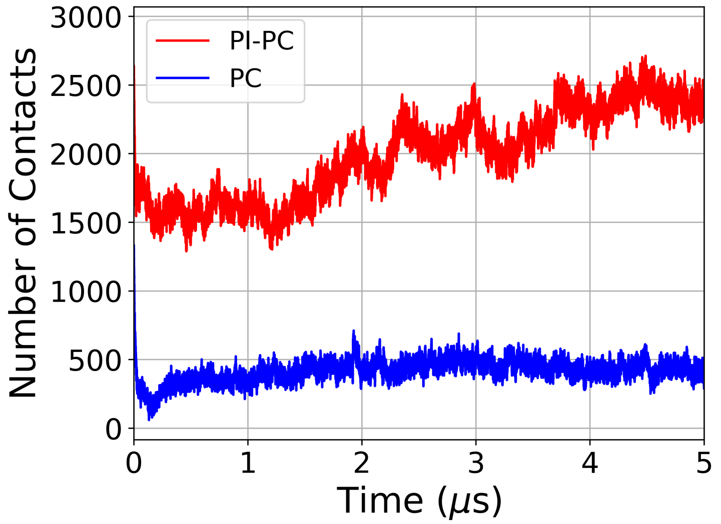

Figure S13: Number of contacts between the tau fibrils and the bilayers for PC–PI and PC systems derived from the coarse grained simulations. The PI infused lipids show more number of contacts. The number of contacts for SF tau with PC systems are included from the coarse grained simulations of our earlier work.

.

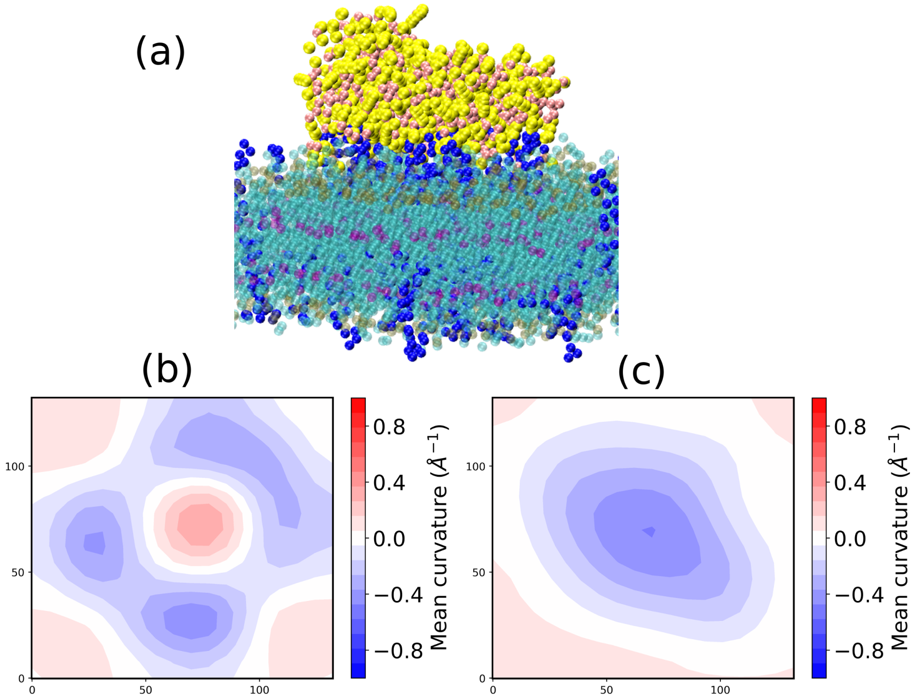

Figure S14: (a) Snapshot of the final configuration of the coarse grained simulations of the tau and the PG–PI lipids. PI lipids are shown in blue using VDW representation. Note the anchoring of the tau fibril by PI lipids. (b)-(c) are the membrane curvature for the PC+PI and PG+PI system respectively. The mean curvature is shown in a colorbar. Negative value of the membrane curvature represents concave surface and positive value of membrane curvature represents convex surface. The mean membrane curvature is calculated using the MDAnalysis python library.

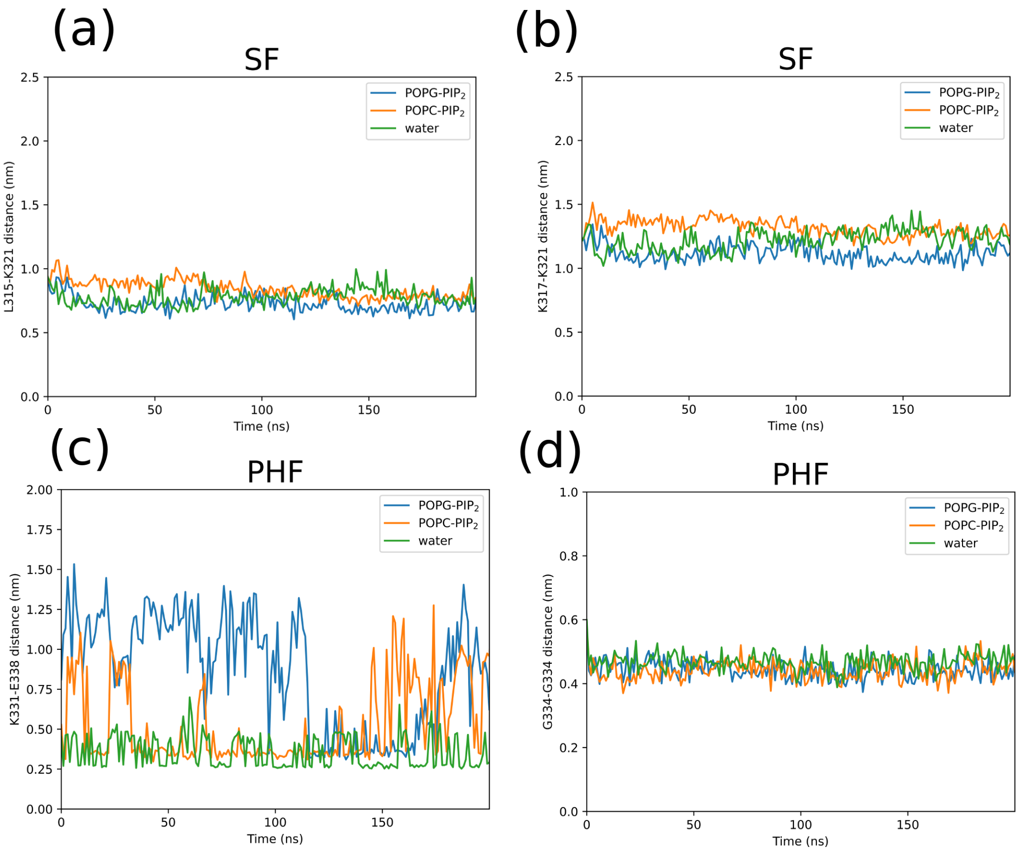

Figure S15: (a)-(b) are the L315–K321 distance and K317–K321 distance in the SF structure. (c)-(d) are the K331–E338 distance and G334–G334 distance in the PHF structure. The residues are shown in the main manuscript.

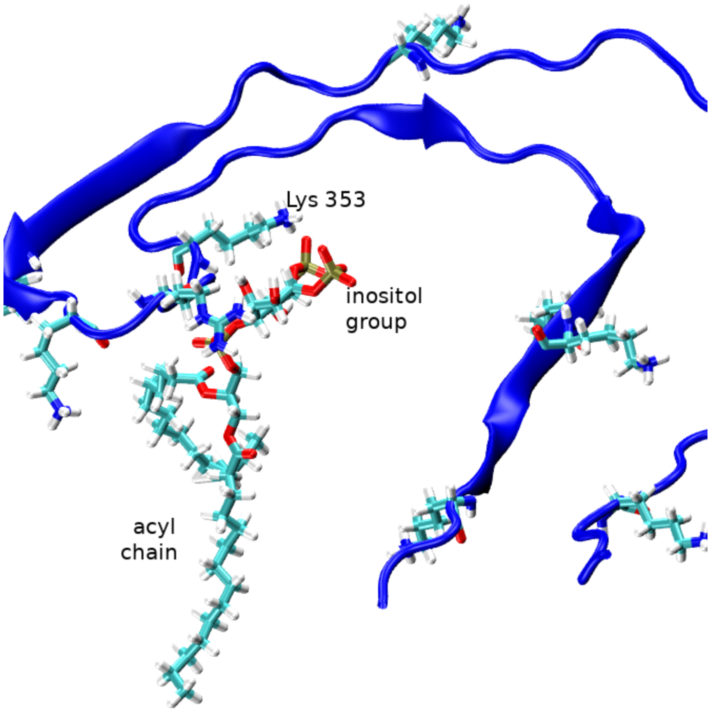

Figure S16: The snapshot of the part of the PHF fibril showing the interaction between the positively charged Lys 353 and the inositol group of the PI lipid in the PC–PI system. The water molecules, lipid molecules and the other chains of the tau fibrils are omitted for clarity.

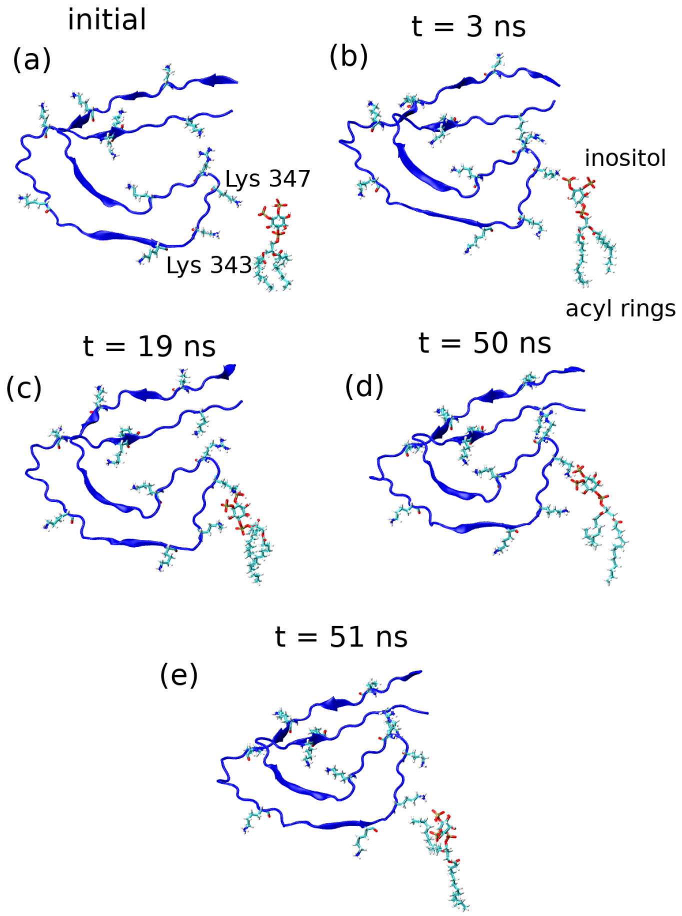

Figure S17: The snapshots of the part of the PHF fibril showing the positive charged residues of tau interacting with the inositol group of PI lipid in the PC–PI system. The water molecules, the other chains of the tau fibril and the PC lipids are omitted for clarity.

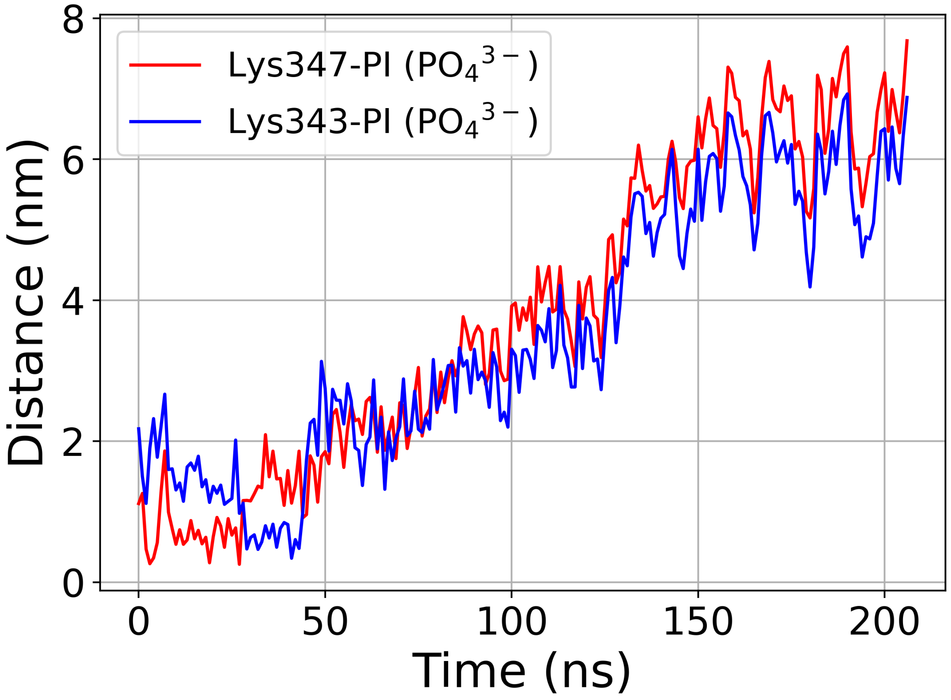

Figure S18: The distance between the phosphate group of the inositol ring of the PI lipid and the residues Lys 343 and Lys 347 of PHF fibril in PC–PI system.
